# Data-driven identification of biomedical systems using multi-scale analysis

**DOI:** 10.1101/2025.06.05.657989

**Authors:** Ismaila Muhammed, Dimitris M. Manias, Dimitris A. Goussis, Haralampos Hatzikirou

## Abstract

Biomedical systems inherently exhibit multi-scale dynamics, making accurate system identification particularly challenging due to the complexity of capturing a wide time scale spectrum. Traditional methods capable of addressing this issue rely on explicit equations, limiting their applicability in cases where only observational data are available. To overcome this limitation, we propose a data-driven framework that integrates the Sparse Identification of Nonlinear Dynamics (SINDy) method, the multi scale analysis algorithm Computational Singular Perturbation (CSP) and neural networks (NNs). This framework allows the partition of the available dataset in subsets characterized by similar dynamics, so that system identification can proceed within these subsets without facing a wide time scale spectrum. Accordingly, when the full dataset does not allow SINDy to identify the proper model, CSP is employed for the generation of subsets of similar dynamics, which are then fed into SINDy. CSP requires the availability of the gradient of the vector field, which is estimated by the NNs. The framework is tested on the Michaelis-Menten model, for which various reduced models in analytic form exist at different parts of the phase space. It is demonstrated that the CSP-based data subsets allow SINDy to identify the proper reduced model in cases where the full dataset does not. In addition, it is demonstrated that the framework succeeds even in the cases where the available data set originates from stochastic versions of the Michaelis-Menten model. This framework is algorithmic, so system identification is not hindered by the dimensions of the dataset.

**Author summary:** Biomedical systems often evolve across multiple time scales, posing major challenges for constructing accurate models directly from data. Traditional model reduction techniques require explicit equations and thus cannot be applied when only observational data are available. To address this, we developed a data-driven framework that combines Sparse Identification of Nonlinear Dynamics (SINDy), Computational Singular Perturbation (CSP) and neural networks (NNs). Our approach automatically partitions a dataset into subsets characterized by similar dynamics, allowing valid reduced models to be identified in each region. When SINDy fails to recover a global model from the full dataset, CSP -leveraging Jacobian estimates from NNs-successfully isolates dynamical regimes where SINDy can be applied locally. We validated this framework using the Michaelis-Menten biochemical model, which is known to admit multiple reduced models in different regions of the phase space. Our method consistently identified the appropriate reduced dynamics, even when the data originated from stochastic simulations. Because our approach is algorithmic and equation-free, it is scalable to high-dimensional systems and robust to noise, offering a promising solution for data-driven model discovery in complex biomedical systems.

## Introduction

Mathematical modeling in biomedical systems often relies on first-principles approaches, typically formulated as differential equations, to describe, predict, and analyze biological processes. Experimental data is also used to calibrate and refine these models [1]. However, the complex, nonlinear, and multi-scale nature of such systems presents significant challenges for deriving accurate governing equations solely through traditional methods [2]. A key difficulty lies in the inability to capture the full spectrum of time scales that characterize the evolution of the system. Model reduction is a major approach to address this limitation, since it combines low dimensionality with the preservation of key dynamical features. Moreover, such reduction enables efficient analysis and interpretation [3]. Various methods have been developed for this purpose, mainly based on available governing equations. Such methods, like Quasi-Equilibrium (QE) [4], Quasi-steady-state Approximation (QSSA) [5], Computational Singular Perturbation (CSP) [6] and the method of invariant manifolds [7–9], decompose the set of variables into fast and slow components, by identifying low-dimensional manifolds and models that govern the long-term dynamics of the slow components.

Mathematical models can also be obtained by extracting dynamics directly from observational data [2]. A variety of data-driven methods have been developed to identify governing equations. These methods employ techniques such as sparsity promotion, symbolic regression, or machine learning to address the multi-scale nonlinear dynamics. Sparse Identification of Nonlinear Dynamics (SINDy) [10] is a prominent method that identifies sparse models by selecting a minimal set of nonlinear functions to capture system dynamics. Weak SINDy [11] improves robustness against noisy and sparse data, especially beneficial for multi-scale systems with variable noise levels. More recently, iNeural SINDy has enhanced this framework by integrating neural networks and using an integral formulation to better handle noisy and sparse datasets [12]. Symbolic regression methods like PySR [13] use evolutionary algorithms to discover closed-form equations, making them suitable for capturing nonlinear behaviors, even in the presence of sparse data. Additionally, Physics-Informed Neural Networks (PINNs) [14] incorporate physical laws into their structure, enabling accurate model predictions from limited data. ARGOS, another symbolic regression method, uses evolutionary algorithms to discover interpretable, sparse models, building on methods like SINDy and PySR while introducing improvements for handling complex systems [15]. Dynamic Mode Decomposition (DMD) [16] and its extended version, EDMD [17], are effective for identifying principal modes and predicting system evolution from relatively sparse data. However, DMD does not recover explicit equations and its accuracy can be limited when the data do not sufficiently capture the underlying dynamics, particularly in multi-scale systems [18].

When the construction of governing equations is impractical or unfeasible, Jacobian estimation methods provide an efficient alternative for analyzing local system behavior. These methods focus on characterizing the local stability or linearization near equilibrium points, which is particularly useful for multi-scale and nonlinear systems. Techniques such as automatic differentiation through NNs [19] approximate the Jacobian matrix from data, allowing the assessment of system sensitivity without requiring a full model. The Lyapunov method, typically used to evaluate equilibrium stability, can also be applied to approximate the Jacobian matrix [20, 21]. Additionally, kernel-based approaches like Gaussian processes estimate Jacobians by fitting smooth functions to data, providing an efficient means to analyze local dynamics in highly nonlinear settings [22]. The Koopman operator theory offers another strategy by linearizing nonlinear systems, enabling finite-dimensional approximations for control and Jacobian estimation [23]. However, data sparsity remains a critical limitation, as sparse sampling, particularly in multi-scale systems, can lead to inaccurate derivative estimates and unreliable Jacobian matrices [18].

Figure 1 provides an overview of the most commonly used data-driven methods for full system identification and Jacobian matrix estimation, categorized by their requirements in terms of the number of variables and data points. Each method has inherent trade-offs, especially when dealing with high-dimensional systems or limited data. As the number of variables increases, the risk of overfitting also rises, particularly in the presence of sparse or noisy data. This challenge is amplified in multi-scale systems, where critical dynamical features span multiple scales, making it difficult to accurately capture system behavior. Neural network-based methods are particularly data-intensive and prone to performance degradation when data is sparse, while other methods that decompose data into modes often require a dense set of observations to capture the full range of dynamics [18, 24]. These challenges underscore the need for hybrid or regularization techniques that can effectively handle multi-scale systems, balancing model complexity with data availability.

**Fig 1.**
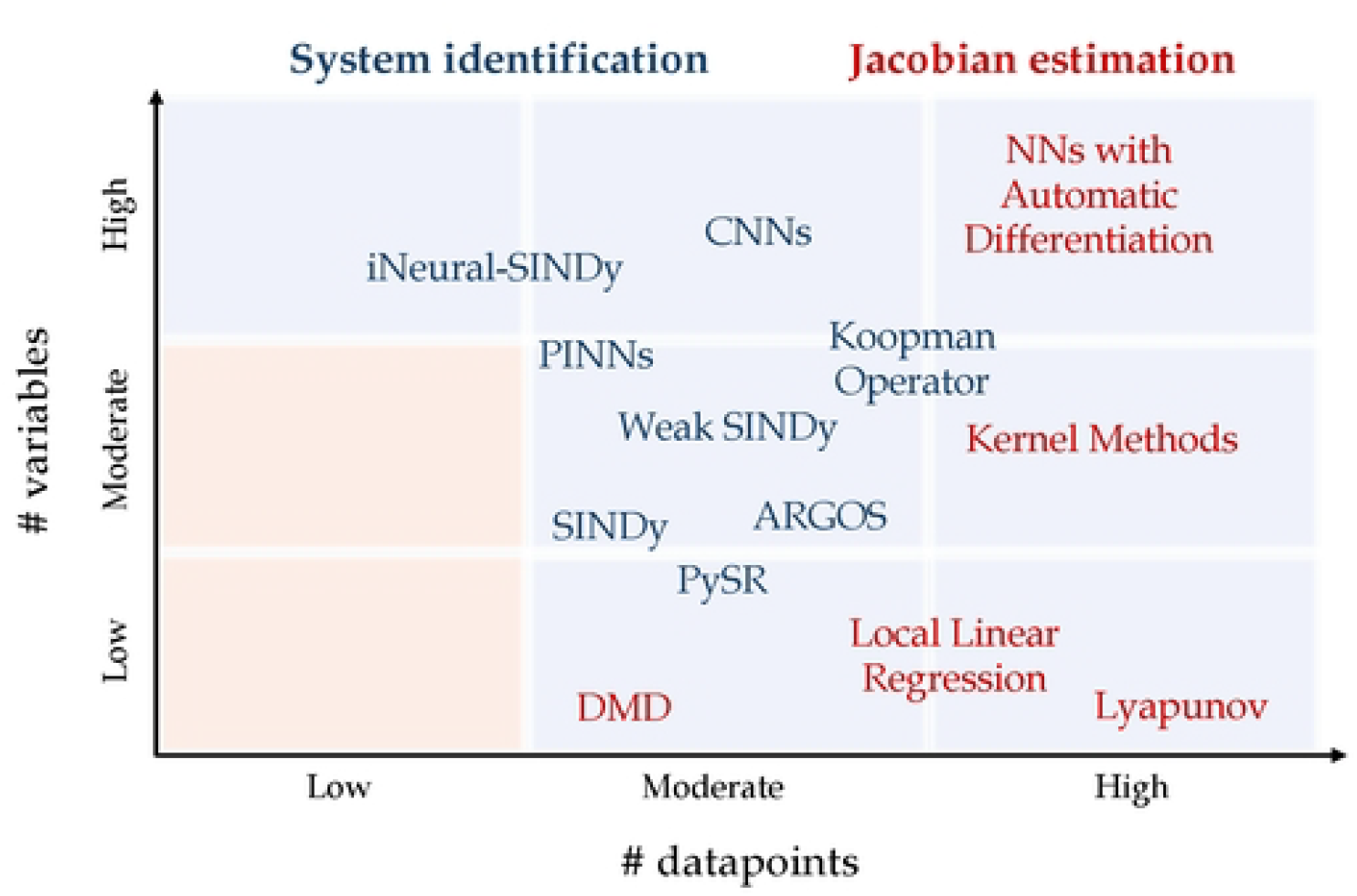
Existing data-driven methods. Schematic representation of existing data-driven methods for full system identification and Jacobian matrix estimation as a function of the number of variables and data points. Each method occupies a distinct region based on its data and dimensionality requirements. No methods can provide accurate estimations in the low variable-low datapoints region.

A novel hybrid framework is introduced here that employs time scale decomposition for model identification in biomedical systems. By integrating the weak formulation of SINDy [10, 11], CSP [25] and NN-based Jacobian estimation [19], the approach identifies algorithmically regions within a given dataset, in which valid reduced models can be constructed. In addition, the approach allows for extracting algorithmically mechanistic insight from multi-scale biological datasets. It is demonstrated that the proposed approach succeeds in cases where existing methodologies fail when applied to the full dataset.

The merits of the proposed framework are demonstrated on the basis of the Michaelis-Menten (MM) model, a widely used and well-established model in biological research for studying enzyme-substrate interactions and reaction kinetics [26, 27]. Due to its simplicity and interpretability, the MM model is extensively used in Systems Biology, biochemistry, and computational biology [28, 29]. Despite its straightforward formulation that allows analytical tractability, the model’s nonlinear interactions and multi-scale dynamics present challenges for conventional system identification, making it a suitable benchmark for evaluating data-driven reduction methods [30, 31].

## Materials and methods

### Sparse Identification of Nonlinear Dynamics (SINDy)

SINDy, introduced by Brunton et. al. [10], is a framework used to identify dynamical systems from time series data using sparse regression. The main idea of SINDy is the assumption that the dynamics of the system can be represented as a sparse combination of candidate functions, making it computationally efficient and interpretable. Weak SINDy, a variant of SINDy was proposed by Schaeffer H. [32], to address the challenges of noise and and irregularly sampled datasets. This method reformulates the system identification problem in a weak form by integrating against a set of test functions, thereby reducing sensitivity to noise and enabling the use of coarsely sampled data. Despite its robustness to noise, the applicability of Weak SINDy is limited to models that conform to predefined functional forms, typically involving linear or weakly nonlinear relationships among variables. It struggles to identify complex models characterized by strong nonlinear interactions or variable-dependent nonlinearity, thereby limiting its effectiveness in systems with intricate dynamical structures [32].

### Data-driven Jacobian Estimation

To perform time scale decomposition and model reduction using CSP, the Jacobian matrix of the system is a key component. When explicit dynamical equations are unavailable, we use a combination of Neural Ordinary Differential Equations (NODE) [33] and NNs [34] to estimate the Jacobian matrix from data. NODE, introduced by Chen et al. [33], provides a way of learning the vector field that governs the continuous-time evolution of a system from data using NNs with the adjoint sensitivity method, which efficiently optimizes the parameters of the system through gradient-based techniques. Frederic et al. introduced a novel approach for training NNs to estimate the Jacobian matrix of an unknown multivariate function using only input-output data pairs (**x, F**(**x**)) [34]. The method utilizes a loss function based on the nearest neighbor search and linear approximations within the sample data.

### The Computational Singular Perturbation Method (CSP)

The CSP algorithm allows for the analysis of multi-scale systems of ordinary differential equations (ODEs), by enabling the local decomposition of the tangent space into fast and slow subspaces [35, 36]. At leading order, the two subspaces can be approximated by the right eigenvectors of the Jacobian, allowing the resolution of the vector field onto fast and slow components [37, 38]. The fast component vanishes over a short transient, implying the equilibration of fast processes and the emergence of constraints that define the *Slow Invariant Manifold* (SIM). On this manifold, the slow dynamics evolve under the influence of the slow component of the vector field [39, 40]. The CSP-based algorithmic vector field decomposition allows system-level identification of dominant fast and slow processes and their influence on the system’s behavior, independent of dimensionality or nonlinearity. CSP has been widely applied to multi-scale systems, such as chemical reaction networks [41–43], biological systems [44, 45] and combustion configurations [46, 47], where time-scale separation presents significant analytical and computational challenges. A mathematical description of the methodology is provided in S1 Appendix.

### Proposed Framework

Figure 2 presents a schematic overview of the proposed framework. Given a time series dataset, SINDy -or any other such method-is first applied, when feasible, to directly identify the governing equations. In cases where SINDy fails, either due to noise, data sparsity or low data volume, NODE [33] is utilized to provide a uniform and dense vector field that is subsiquently used in a NN [34] to estimate the Jacobian matrix. Then, the estimated Jacobian matrix is considered for CSP to be applied without the need for explicit governing equations. The eigenvectors and eigenvalues of the estimated Jacobian serve as leading-order approximations of the system’s corresponding fast/slow directions and time scales. Using the CSP theory and diagnostic tools [44, 48, 49], the dataset is then analyzed to identify regions where valid models can be constructed. These regions enable the partitioning of the dataset into subsets, each corresponding to a different dynamical regime. Finally, SINDy is applied to each subset independently to derive region-specific models that accurately capture the dynamics within each regime.

**Fig 2.**
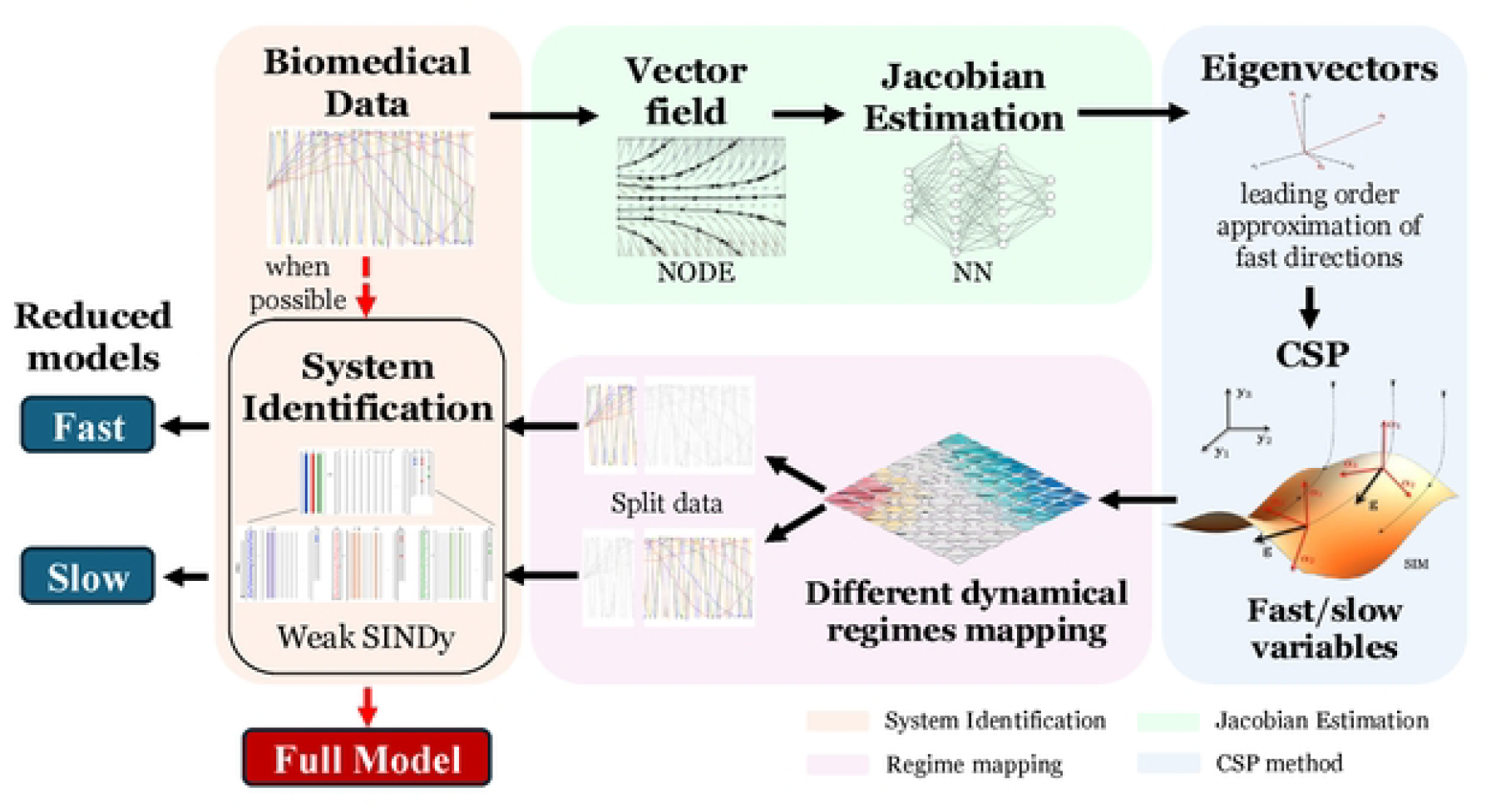
Framework. The methodology proposed to identify regions for valid models when system identification methodologies fail.

## Results

We demonstrate the application of our methodology to the multi-scale Michaelis-Menten model, which can be simplified to three different reduced models that are valid in tdifferent domains of the phase and parameter spaces; the standard Quasi-Steady-State Approximation (sQSSA), the reverse Quasi-Steady-State Approximation (rQSSA) and the Partial Equilibrium Approximation (PEA) [31, 45]. Although the model was introduced more than a century ago [26], its popularity keeps increasing [50] due to its significance in many biological and medical contexts [51].

### The Reduced Michaelis-Menten Models

#### Equation-based Model Reduction

The MM reaction mechanism describes the interaction between a substrate *S* and an enzyme *E*, forming a reversible complex *C*. The complex *C* subsequently undergoes an irreversible reaction, yielding a product *P* and releasing the enzyme *E*, which can then participate in another cycle of substrate binding [26]. This process can be represented as follows:

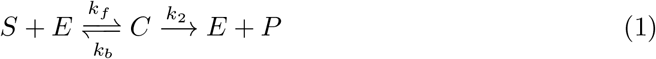

where *k*_*f*_ (L mol^−1^*s*^−1^) and *k*_*b*_ (*s*^−1^) denote the forward and reverse rate constants of the enzyme-substrate complex formation, respectively, while *k*_2_ (*s*^−1^) represents the catalytic rate constant, also referred to as the turnover number. According to the law of mass action, the MM mechanism is modeled by the following set of ordinary differential equations (ODEs):

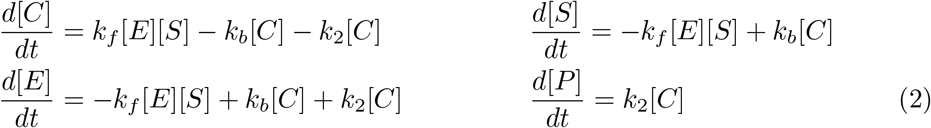

Here, the square brackets denote the concentrations of the respective chemical species. Assuming the system is closed, where initially no product or complex is present, i.e., [*C*](0) = [*P*](0) = 0, the conservation relations [*E*] + [*C*] = [*E*](0) and [*S*] + [*C*] + [*P*] = [*S*](0) hold ([*E*](0) and [*S*](*O*) are the initial enzyme and substrate concentrations), the system in Eq. (2) simplifies to:

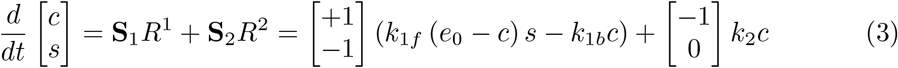

where the simplified symbols *c* = [*C*], *s* = [*S*] and *e*_0_ = [*E*](0) have been used. Here **S**_1_ and **S**_2_ are the stoichiometric vectors of the two reactions and *R*^1^ = *R*^1*f*^ − *R*^1*b*^ = *k*_1*f*_ (*e*_0_ − *c*)*s* − *k*_1*b*_*c* and *R*^2^ = *k*_2_*c* are the related rates. It should be noted that the origin, (*c, s*) = (0, 0), is the only equilibrium point of the system.

The multi-scale nature of the two-dimensional system in Eq. (3) is manifested through the significant difference in magnitude between the two time scales, *τ*_1_ and *τ*_2_ (*τ*_1_ *< τ*_2_), which govern the system’s dynamics. These time scales can be approximated by the inverse modulo of the eigenvalues *λ*_+_ and *λ*_−_ of the Jacobian matrix *J* of the two-dimensional ODE system described in Eq. (3) [35, 42, 52]:

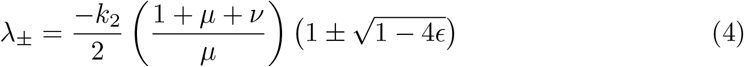

where

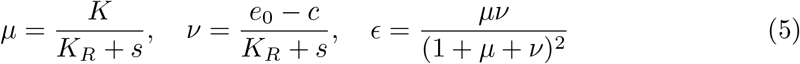

and *K*_*R*_ = *k*_1*b*_*/k*_1*f*_ is the dissociation constant, *K* = *k*_2_*/k*_1*f*_ is the Van Slyke-Culen constant and *K*_*M*_ = *K*_*R*_ + *K* is the Michaelis-Menten constant [31]. The value of *ϵ* is indicative to the gap that develops between the two eigenvalues (and corresponding time scales) and defines the stiffness of the system; i.e., *τ*_1_*/τ*_2_ = *f*(*ϵ*), so that *τ*_1_*/τ*_2_ *≪* 1 when *ϵ ≪*1 and *τ*_1_*/τ*_2_ *→*1 when *ϵ →* 1*/*4 [53, 54]. Both *µ* and *ν* are non-dimensional and non-negative and quantify the relative availability and initial distribution of enzyme and substrate in dimensionless form, determining the model’s dynamic behavior and the separation between fast and slow reaction processes. The analytical expression of the eigenvalues in terms of *µ* and *ν* allows us identify regions in the *µ*-*ν* plane where different reduced models can be constructed [55].

Figure 3 displays the regions of validity of different reduced models. In particular, the region of a valid sQSSA model is highlighted with pink, the region of a valid rQSSA model is highlighted with green and the region of valid Partial Equilibrium Approximation (PEA) model is highlighted with shaded blue lines. The regions are separated by the lines *ν* = 1 + *µ* and *µ* = 1 + *ν*. Along the *µ <* 1 narrow neighborhood of the *ν* = 1 + *µ* line (solid), indicated by fading to white color, no valid QSSA model can be obtained and only the PEA model is valid. Along the *µ >* 1 part of the *ν* = 1 + *µ* line (solid) *ϵ →* 1*/*4, so that no reduced model can be constructed there since *τ*_1_*/τ*_2_ *→* 1. Moving away from this part, *ϵ* progressively decreases, as it is indicated by the *ϵ* = 10^−2^ dashed curve [31, 55]. Table 1 displays the reduced models that are valid in the specified portions of the *µ* − *ν* plane; i.e., the sQSSA model (*c* is considered fast), the rQSSA model (*s* is considered fast) and the PEA model (the bi-directional reaction *S* + *E ? C* is in equilibrium, *R*^1*f*^ ≈*R*^1*b*^). It is shown in Table 1 that the PEA model simplifies to the sQSSA model when *ν ≪*1 and to the rQSSA model when *ν ≫*1.

**Fig 3.**
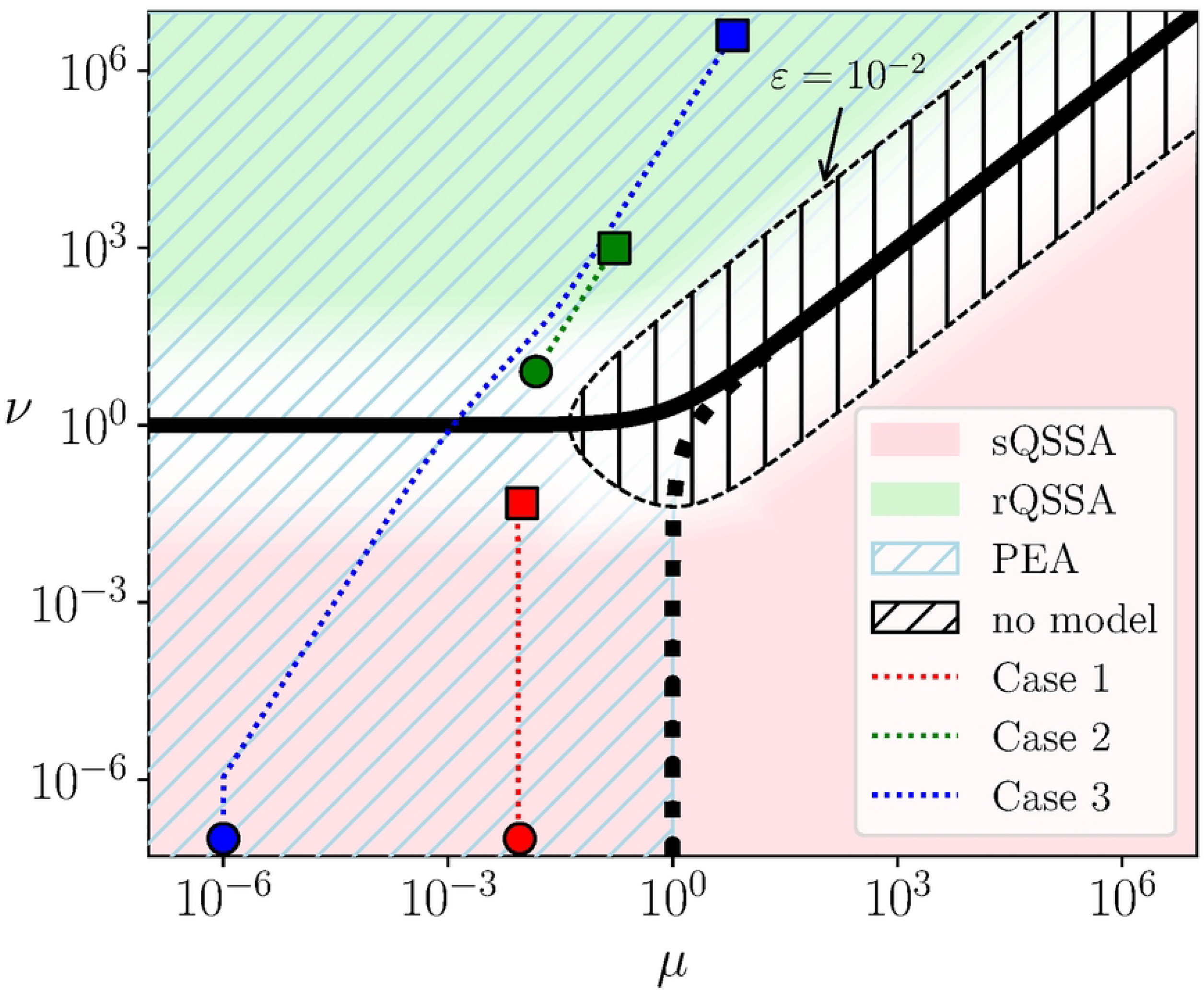
Regions of valid reduced models in the *µ*-*ν* plane. The regions of validity of sQSSA (pink: *ν ≪µ* + 1), rQSSA (green : *ν ≫ µ* + 1) and PEA (shaded blue: *µ≪ν* + 1) in the *µ*-*ν* plane, along with the trajectories that are analyzed (see Table 2 for the related parameters and ICs). Circles and squares denote the starting and ending point of each trajectory, respectively. The thick solid and dotted lines denote *ν* = 1 + *µ* and *µ* = 1 + *ν*, respectively. The thin dashed curve denotes the points at which *ϵ* = 10^−2^ and encapsulates the shaded region where *ϵ >* 10^−2^. Reduced models in this region are of low accuracy [55].

The fact that the full model in Eq. (3) is valid throughout the *µ* − *ν* plane and the three reduced models in Table 1 are valid in portions of this plane, will be the basis for the assessment of the proposed framework for system identification. In particular, datasets originating from the three trajectories shown in Fig. 3 will form the starting point of the identification process. As shown in the figure one trajectory is located in the region where the sQSSA and the PEA models are valid (Case 1), another is located in the region where the rQSSA and the PEA models are valid (Case 2) and a third trajectory is located in a region that includes the domains of validity of the sQSSA/PEA models (first part of the trajectory), the PEA model (middle part) and the rQSSA/PEA models (last part) (Case 3). The parameters and initial conditions for the three trajectories are displayed in Table 2.

**Table 1.** The algebraic relation that defines the SIM and the differential equation(s) that governs the flow on the SIM, according to sQSSA, rQSSA and PEA.

| Region | Algebraic Eq. (SIM) | Differential Eq. (reduced system) |
| --- | --- | --- |
| sQSSA | $(e_0 - c)s - K_R c - Kc = 0$<br>$\left[ c = \frac{e_0 s}{K_M + s} \right]$ | $\frac{ds}{dt} = -k_2 c$<br>$\left[ \frac{dc}{dt} = -\frac{k_2 (e_0 - c)^2 c}{K_M e_0} \right]$ |
| rQSSA | $(e_0 - c)s - K_R c = 0$<br>$\left[ s = \frac{K_R c}{e_0 - c} \right]$ | $\frac{dc}{dt} = -k_2 c$<br>$\left[ \frac{ds}{dt} = -\frac{k_2 (K_R + s)s}{K_R} \right]$ |
| PEA | $(e_0 - c)s - K_R c - \frac{1}{1+\nu} Kc = 0$ | $\frac{dc}{dt} = -\frac{\nu}{1+\nu} k_2 c$<br>$\frac{ds}{dt} = -\frac{1}{1+\nu} k_2 c$ |

In brackets are the traditional forms of the relations that approximate the SIM and the differential equation that governs the evolution of the fast variable in the sQSSA and rQSSA context.

**Table 2.** Parameters and initial conditions for simulation of the cases considered.

| Case | $k_{1f}$ | $k_{1b}$ | $k_2$ | $e_0$ | $s(0)$ | $c(0)$ |
| --- | --- | --- | --- | --- | --- | --- |
| Case 1 | 10 | 100 | 1 | 0.5 | 1 | 0.045045 |
| Case 2 | 3 | 3 | 0.5 | 1000 | 10.11111 | 910 |
| Case 3 | 1000 | 0.01 | 0.1 | 10 | 10 | 9.999999 |

The choice of the values is based on the analysis in [56] and [55]. Initial conditions are obtained on the *SIM* for each case.

The validity of the reduced models in Table 1 is demonstrated in Fig.4, where profiles of the three trajectories considered here obtained with the full and appropriate reduced model are compared. It is shown that an excellent agreement is obtained.

The simplification of the PEA model to the two QSSA models in Case 3 is demonstrated in Fig. 5 that compares the solution of the PEA model with the solutions provided by the two QSSA models of Table 1. The sQSSA-based profiles of *c* and *s* in the left panel and the rQSSA profiles in the right panel are denoted by dashed lines, while the PEA profiles in both panels are denoted by crosses. It is evident that the sQSSA solution closely follows the original trajectory in the first region (pink, as in Fig 3), where the sQSSA is valid, but begins to diverge as the system transitions into the second region (green, as in Fig 3), where only rQSSA is valid. Conversely, the rQSSA solution initially deviates from the original trajectory in the first region, where rQSSA is not valid, but progressively aligns with it upon entering the second region, where rQSSA is valid.

### Data-driven Reduced Model Identification using SINDy

In order to assess the validity of the proposed method for system identification, datasets were produced from the profiles shown in Fig. 4 from both the full and reduced models, by considering a relatively sparse and uniform grid of *n*=100 datapoints; a limit close to where SIDNy might fail.

**Fig 4.**
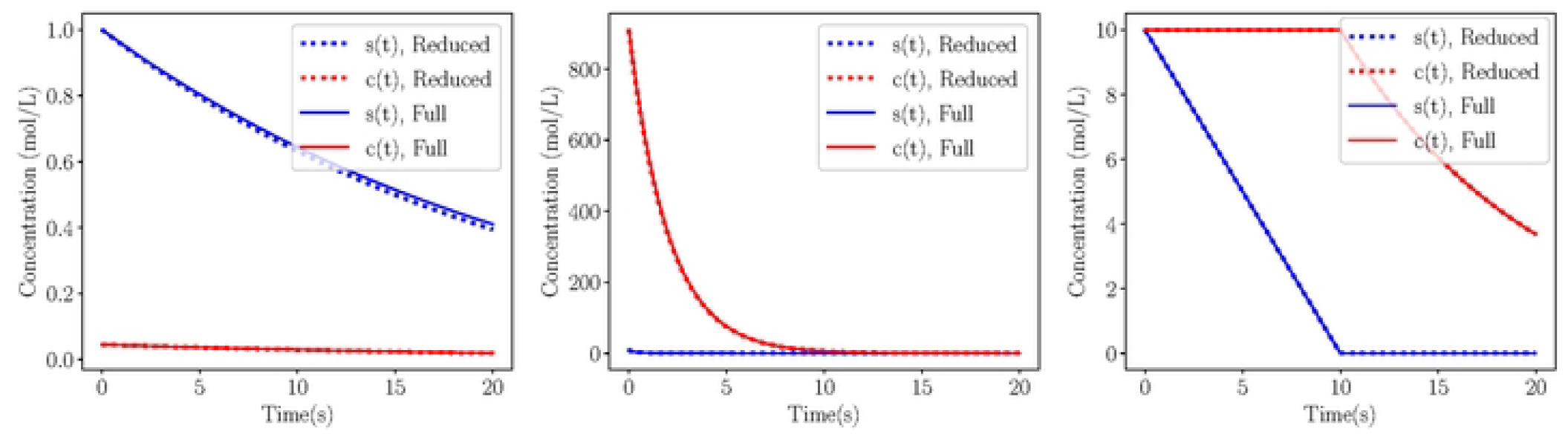
System Solutions. Solution comparison between full (solid) and reduced (dotted) Michaelis-Menten simulations on the SIM, for Case 1 - sQSSA (left), Case 2 - rQSSA (middle) and Case 3 - PEA (right). Parameters and initial conditions on Table 2.

**Fig 5.**
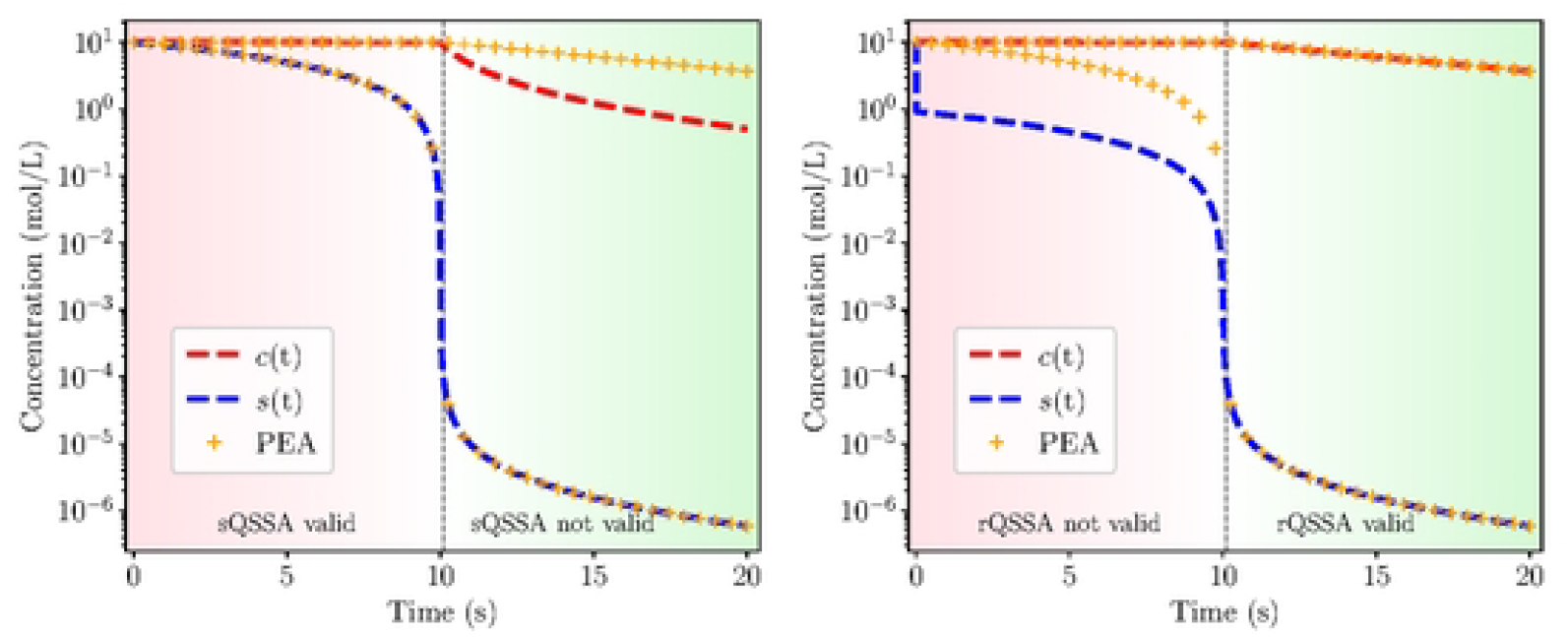
Validity comparison. Validity comparison between the sQSSA (left) and rQSSA (right) and PEA solutions of Case 3 (see Table 2 for parameters and ICs).

The solutions of each case, both of the full and the reduced models, were used as datasets in Weak SINDy for system identification. The results are displayed in Table 3. The *coefficient of determination*, denoted as *R*^2^, was used to evaluate how well the identified models fit the data. *R*^2^ measures the proportion of the variance in the dependent variable (typically the system’s dynamics) that is predictable from the independent variables (the terms in the identified governing equations).

The coefficient of determination is computed as:

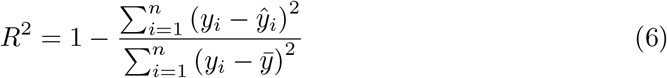

where *y*_*i*_ indicates the actual observed data (here, the derivative of the state,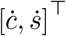) and *ŷ*_*i*_ indicates the predicted values from the SINDy model. The mean obsearved data is given by 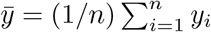. When *R*^2^ = 1, the predicted model explains all the variance in the data.

The results in Table 3 indicate that Weak SINDy successfully identified the different models corresponding to the data generated from the full models. In particular, it accurately recovered the parameters of each system, as reflected by the high coefficient of determination values (*R*^2^), which are consistently close to unity.

When applied to data from the reduced models, Weak SINDy accurately identified the reduced dynamics in the two QSSA cases. However, in Case 3 the method failed to recover the reduced model from the data. Instead, it returned a polynomial approximation resembling the full model. This discrepancy is reflected in the large negative values of *R*^2^, indicating that the identified model structure does not capture the underlying dynamics accurately.

This failure is attributed to the limitations of SINDy in constructing models with complex nonlinear terms that do not conform to its predefined candidate library. The

The coefficient of determination (*R*^2^) indicates the accuracy of the Weak SINDy reconstruction.

PEA reduced model includes nonlinear terms of the form 1*/*(1 + *ν*) and *ν/*(1 + *ν*), where *ν* = (*e*_0_ −*c*)*/*(*K*_*R*_ + *s*) is itself a nonlinear function of the state variables. Since SINDy relies on a fixed library of basis functions, typically comprising polynomials, trigonometric functions, or other simple expressions, it struggles to represent such composite nonlinearities. However, when SINDy is directly provided with functions that explicitly include the nonlinear terms of the numerators and denominators, it successfully captures the dynamics and correctly identifies the PEA reduced model [10]. Despite this, it remains impractical to know a priori the specific nonlinear terms required for accurate model identification in real-world applications, where the underlying functional forms are typically unknown. This highlights a fundamental challenge when using SINDy for complex multi-scale systems with non-standard interactions.

**Table 3.** Comparison of the right-hand side expressions between the ground truth and the identified models, using data obtained from the deterministic full (top) and reduced (bottom) models.

| | Case | Ground Truth | Identified Model | $R^2$ |
| --- | --- | --- | --- | --- |
| Full Model | Case 1 | $\dot{c} = 5s - 101c - 10sc$ | $\dot{c} = 5.004s - 101.073c - 10.003sc$ | 0.9991 |
| | | $\dot{s} = -5s + 100c + 10sc$ | $\dot{s} = -5.003s + 100.07c + 10.003sc$ | 0.9990 |
| | Case 2 | $\dot{c} = 3000s - 3.5c - 3sc$ | $\dot{c} = 2999.915s - 3.5c - 3sc$ | 1.0 |
| | | $\dot{s} = -3000s + 3c + 3sc$ | $\dot{s} = -3000.007s + 3c + 3sc$ | 1.0 |
| | Case 3 | $\dot{c} = 100000s - 0.11c - 10000sc$ | $\dot{c} = -1000004.014s - 0.11c - 10000.401sc$ | 0.9959 |
| | | $\dot{s} = -100000s + 0.01c + 10000sc$ | $\dot{s} = -100004.038s + 0.01c + 10000.404sc$ | 0.9975 |
| Reduced Model | Case 1 | $\dot{c} = -0.050cc + 0.198c^2 - 0.198c^3$ | $\dot{c} = -0.050c + 0.198c^2 - 0.198c^3$ | 0.9999 |
| | | $\dot{s} = -1c$ | $\dot{s} = -0.999c$ | 1.0 |
| | Case 2 | $\dot{c} = -0.5c$ | $\dot{c} = -0.5c$ | 0.9996 |
| | | $\dot{s} = -0.5s - 0.5s^2$ | $\dot{s} = -0.5s - 0.5s^2$ | 1.0 |
| | Case 3 | $\dot{c} = -\frac{\nu}{1+\nu}0.1c$ | $\dot{c} = -5944980.763s + 0.620c + 594498.076sc$ | -2.5614 |
| | | $\dot{s} = -\frac{1}{1+\nu}0.1c$ | $\dot{s} = 5945000.197s - 0.720c - 594500sc$ | -4.3303 |

The coefficient of determination ( $R^2$ ) indicates the accuracy of the Weak SINDy reconstruction.

### Implementing our proposed method for Case 3

The framework proposed here addresses the challenges encountered by Weak SINDy in identifying a valid reduced model in Case 3 as follows. First, the NODE network [33] is employed to generate a dense and uniform vector field, needed for the accurate implementation of the NN-based Jacobian approximation. NODE is utilized in an unsupervised manner, as the objective is to infer the underlying dynamical system from the observed trajectories without explicit labels. Subsequently, the NN methodology proposed in [34] is used to estimate the structure of the Jacobian matrix directly from the reconstructed vector field values, without requiring additional supervision. Hyperparameters, such as nearest neighbors, neighborhood radius and NN architecture, as well as training settings including optimizer and training parameters, are carefully tuned to ensure accurate and stable Jacobian estimation.

The eigenvectors of the Jacobian matrix are then employed by CSP to approximate the fast and slow directions in phase space. Additionally, the inverse of the eigenvalue moduli are utilized to approximate the characteristic time scales of the system; see Eq. (13). When providing a full dataset produced from the PEA model for Case 3, CSP analysis revealed that the dataset contains two distinct regions where vastly different dynamics prevail. In agreement to the results in Fig. 3, CSP concluded that: (i) in the first region, the enzyme-substrate complex *c* exhibits fast dynamics, justifying the applicability of sQSSA; and (ii) in the second region, the substrate *s* transitions to the fast variable, making rQSSA a valid approximation. Furthermore, CSP explicitly identified the narrow region, where the transition between these two dynamical regimes is realized, allowing for an appropriate partitioning of the dataset in two parts, so that the data in each set are characterized by similar dynamics. Details of the CSP analysis are presented in S2 Appendix.

Following this dataset partitioning, each subset corresponding to a different QSSA model is independently processed using the Weak SINDy algorithm. The results, summarized in Table 4, demonstrate that Weak SINDy successfully identified the appropriate QSSA model within each region.^1^ A comparison between the PEA and identified models in Table 4 might suggest that the zero derivatives of the fast variables are incorrect. However, a closer inspection reveals that these results are structurally consistent with the expected reduced dynamics. This is shown in Fig. 6, which illustrates the temporal evolution of the terms 1*/*(1 + *ν*) and *ν/*(1 + *ν*) in the PEA model over the full data set *ν* ≈ 0, leading to. In the first period, which refers to the data subset where sQSSA is valid, Fig. 6 shows that 1*/*(1 + *ν*) ≈1 and *ν/*(1 + *ν*) ≈0. In these limits, the PEA model simplifies to the identified model. In the second period, which refers to the data subset where rQSSA is valid, 1*/*(1 + *ν*) ≈ 0 and *ν/*(1 + *ν*) ≈ 1. Again, in these limits the PEA model simplifies to the identified model.

**Table 4.** Comparison of the right-hand side expressions between the PEA and the identified models by the Weak SINDy, using the CSP-based split of the dataset of Case 3.

| Region | | PEA model | Identified model | $R^2$ |
| --- | --- | --- | --- | --- |
| Case 3 | Split 1 (sQSSA) | $\dot{c} = -\frac{\nu}{1+\nu} 0.1c$ | $\dot{c} = 0$ | -1.1969 |
| | | $\dot{s} = -\frac{1}{1+\nu} 0.1c$ | $\dot{s} = -0.1c$ | 0.9999 |
| | Split 2 (rQSSA) | $\dot{c} = -\frac{\nu}{1+\nu} 0.1c$ | $\dot{c} = -0.1c$ | 1.0 |
| | | $\dot{s} = -\frac{1}{1+\nu} 0.1c$ | $\dot{s} = 0$ | -7.7547 |

Data obtained from the PEA model. $R^2$ indicates coefficient of determination of Weak SINDy.

**Fig 6.**
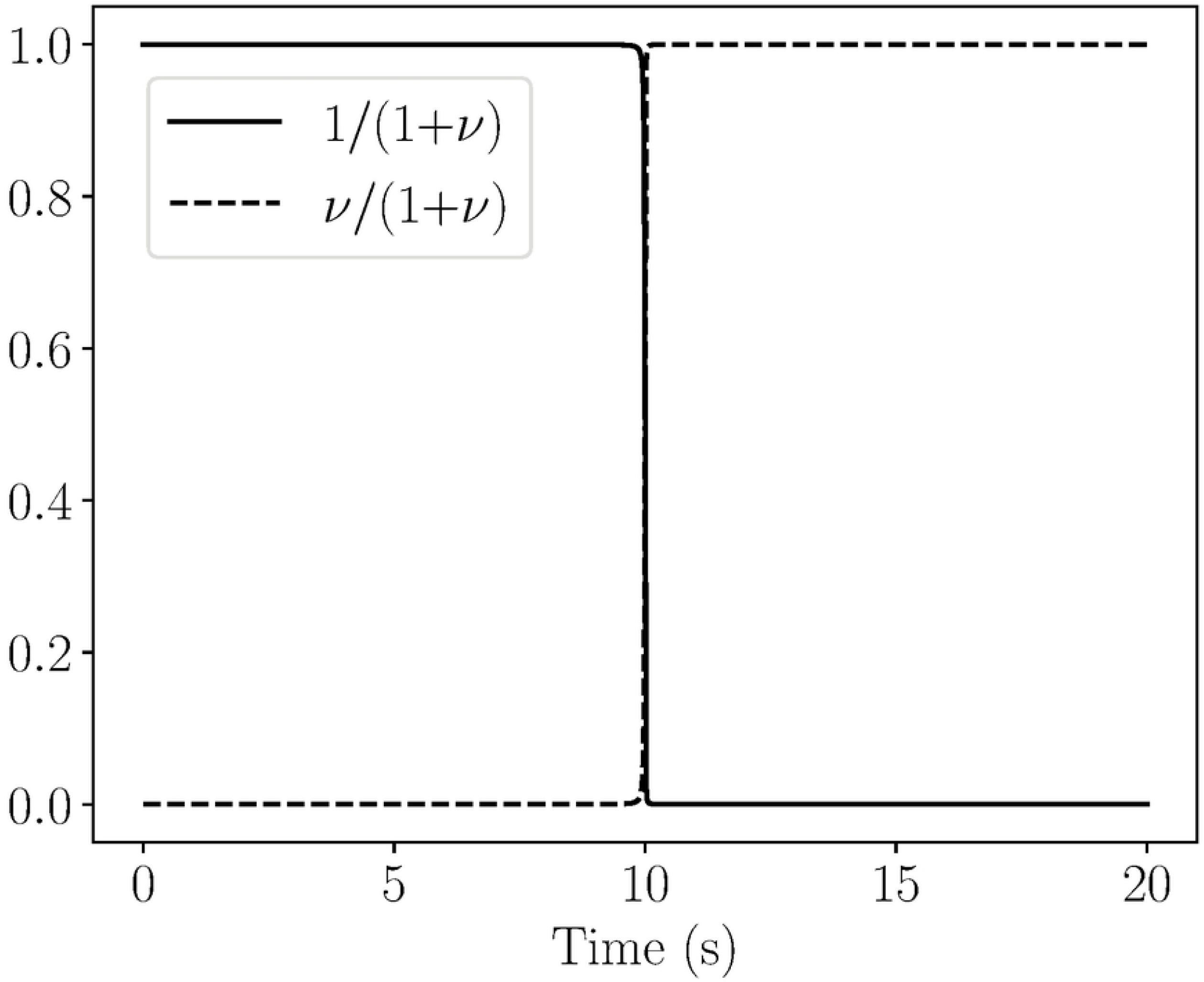
Evolution of 1/(1 + *ν*) and *ν*/(1 + *ν*). Evolution of 1*/*(1 + *ν*) (solid) and *ν/*(1 + *ν*) (dashed) for Case 3. In the first part where sQSSA is valid, 1*/*(1 + *ν*) ≈ 1 and *ν/*(1 + *ν*) ≈ 0, while in the second part where rQSSA is valid, 1*/*(1 + *ν*) ≈ 0 and *ν/*(1 + *ν*) ≈ 1.

The negative values of *R*^2^ in Table 4 arise when the derivatives of the fast variables are set to zero. These negative values do not indicate incorrect dynamics; rather, they reflect limitations of the *R*^2^ metric when applied to variables with negligible variance, as previously discussed. Conversely, the negative *R*^2^ values observed in Table 3 (PEA case) represent genuine model mismatch, since Weak SINDy returns a polynomial approximation inconsistent with the nonlinearities inherent to the PEA reduced dynamics. Thus, while negative *R*^2^ values in Table 3 indicate limitations of Weak SINDy in capturing certain nonlinear forms, those in Table 4 are artifacts of the metric in low-variance contexts.

It was demonstrated here that, while Weak SINDy initially failed to identify a valid global model for the entire dataset, the proposed framework successfully partitioned the data into dynamically distinct regions. Within these regions, Weak SINDy accurately recovered locally valid reduced models, emphasizing that the primary objective in a purely data-driven setting is to identify models that capture local dynamics rather than a single globally valid formulation.

### Model Identification For Noisy Data

In real-world biomedical applications, data contain inherent noise and may exhibit underlying time scale separation. This further complicates the identification of governing dynamics and the construction of valid models using methods such as SINDy, and even its robust variant, Weak SINDy. To simulate noise in the dynamics, we modify the deterministic system in Eq. (3) by introducing stochastic perturbations in two forms: additive and multiplicative noise.

Additive noise is introduced by adding a distrurbance in the right-hand side of the system with Gaussian noise independent of the vector field:

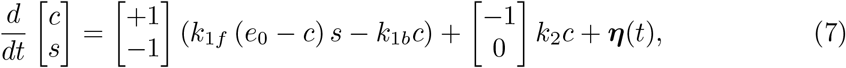

where ***η***(*t*) = [*η*_*c*_(*t*), *η*_*s*_(*t*)]^*⊤*^ is a vector-valued Gaussian noise process with zero mean and standard deviation proportional to the signal magnitude:

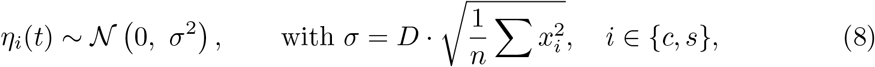

*x*_*i*_ denotes the value of the respective variable at time *t* and *D* denotes the noise percent or strength.

Multiplicative noise is modeled as a stochastic perturbation that scales with the magnitude of the vector field:

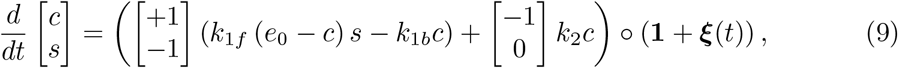

where denotes element-wise multiplication, **1** is a vector of ones matching the state dimension and ***ξ***(*t*) = [*ξ*_*c*_(*t*), *ξ*_*s*_(*t*)]^*⊤*^ is a vector-valued Gaussian noise process with zero mean and standard deviation proportional to the signal magnitude:

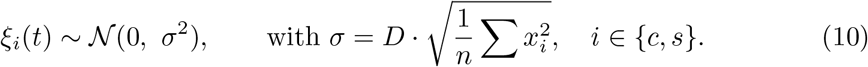

In both cases, noise is introduced at the level of the vector field (i.e., the right-hand side of the ODEs), simulating realistic measurement or process noise often observed in biomedical data. Noisy datasets were generated for both the full and reduced models, derived from Eq. (3) and Table 1, respectively.

The results obtained by applying Weak SINDy to the noisy datasets are summarized in Table 5. For Cases 1 and 2, Weak SINDy successfully identified reduced models, with the right-hand side of the fast variable equations correctly approximated as zero, consistent with the quasi-steady-state assumption. In contrast, the method failed to reconstruct any valid model for Case 3, which involves more complex multi-scale dynamics, under noisy conditions. Figure 7 illustrates the impact of additive (2%) and multiplicative (1%) noise on the system trajectories in Case 3. While additive noise did not significantly alter the trajectory, multiplicative noise led to substantial deviations from deterministic values, especially at higher magnitudes, preventing Weak SINDy from correctly identifying the underlying dynamics. This limitation underscores the challenge posed by the combined effects of multi-scale behavior and noise, emphasizing the necessity of employing the proposed framework for more robust model identification under realistic, noisy biomedical data scenarios.

**Table 5.**
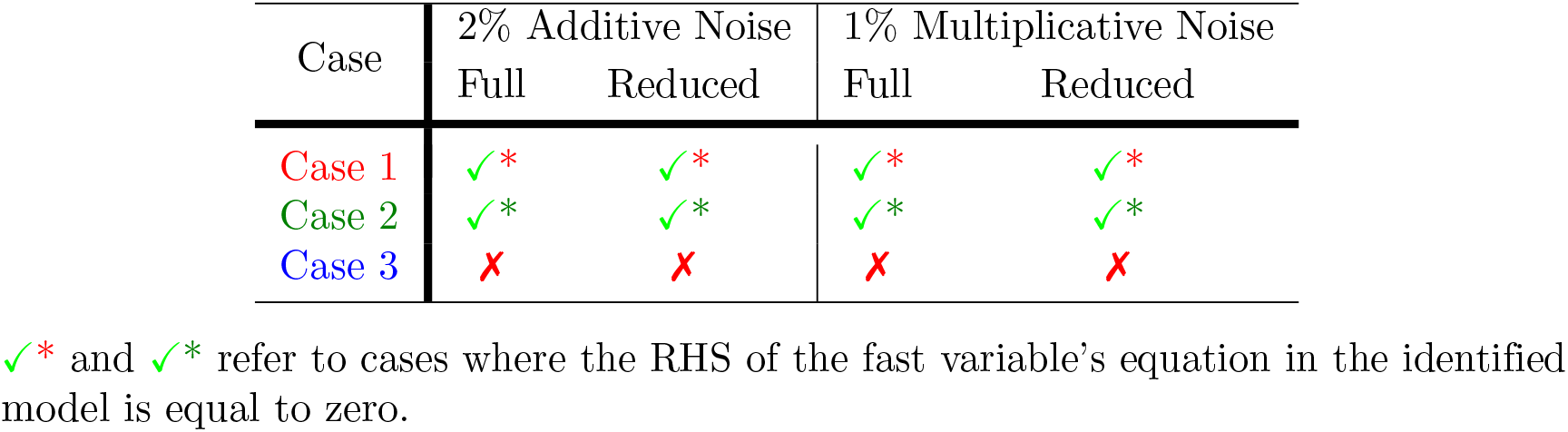
Performance of Weak SINDy on identifying the corresponding models of the data produced from the full and reduced stochastic models.

**Fig 7.**
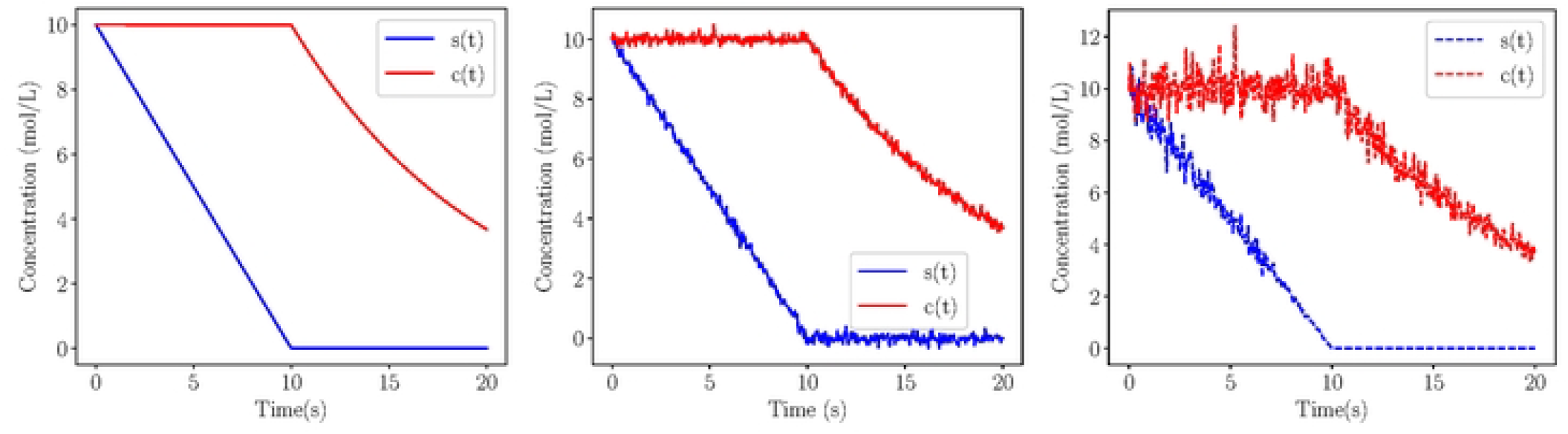
System evolution under noisy conditions. Temporal profiles of the variables reconstructed from NODE for the PEA case: deterministic data (left), data with 2% additive noise (middle) and data with 1% multiplicative noise (right).

Table 6 presents the results of applying Weak SINDy to both deterministic and noisy datasets of Case 3. In all scenarios, the method identified structurally valid reduced QSSA models, correctly setting the right-hand side of the fast variable to zero. Although the associated *R*^2^ values are negative, this outcome stems from the vanishing variance of fast variable derivatives under QSSA assumptions, which limits the metric’s reliability in such contexts. The consistent recovery of valid models across varying noise levels underscores the robustness of the proposed framework.

**Table 6.**
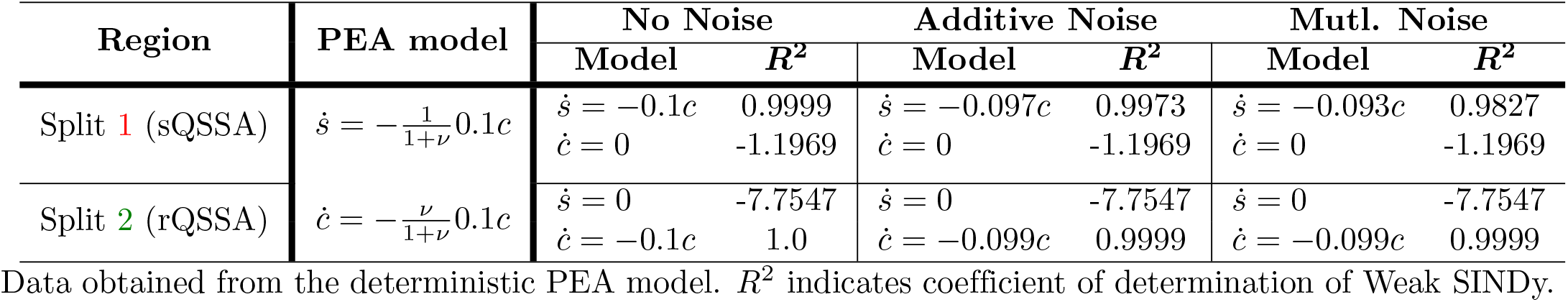
Comparison of the right-hand side expressions between the PEA and the identified models, using the CSP-based split of the dataset of Case 3.

| Region | PEA model | No Noise |  | Additive Noise |  | Mutl. Noise |  |
| --- | --- | --- | --- | --- | --- | --- | --- |
| | | Model | $R^2$ | Model | $R^2$ | Model | $R^2$ |
| Split 1 (sQSSA) | $\dot{s} = -\frac{1}{1+\nu}0.1c$ | $\dot{s} = -0.1c$ | 0.9999 | $\dot{s} = -0.097c$ | 0.9973 | $\dot{s} = -0.093c$ | 0.9827 |
| | | $\dot{c} = 0$ | -1.1969 | $\dot{c} = 0$ | -1.1969 | $\dot{c} = 0$ | -1.1969 |
| Split 2 (rQSSA) | $\dot{c} = -\frac{\nu}{1+\nu}0.1c$ | $\dot{s} = 0$ | -7.7547 | $\dot{s} = 0$ | -7.7547 | $\dot{s} = 0$ | -7.7547 |
| | | $\dot{c} = -0.1c$ | 1.0 | $\dot{c} = -0.099c$ | 0.9999 | $\dot{c} = -0.099c$ | 0.9999 |

Data obtained from the deterministic PEA model. $R^2$ indicates coefficient of determination of Weak SINDy.

Figure 8 compares the deterministic solutions (solid lines) of the PEA model to those identified by Weak SINDy (dotted lines) after partitioning the dataset into the two subsets in which sQSSA (top row) and rQSSA (bottom row) are valid. The three columns correspond to no noise (left), additive noise (middle), and multiplicative noise (right). All data were smoothed using the NODE framework to enable accurate vector field reconstruction. Across all noise levels, the identified trajectories for substrate s(t) (blue) and complex c(t) (red) align closely with the PEA trajectories, demonstrating the effectiveness of the framework in identifying valid reduced models even in the presence of noise. The results of the CSP analysis of the stochastic cases are presented in S3 Appendix.

**Fig 8.**
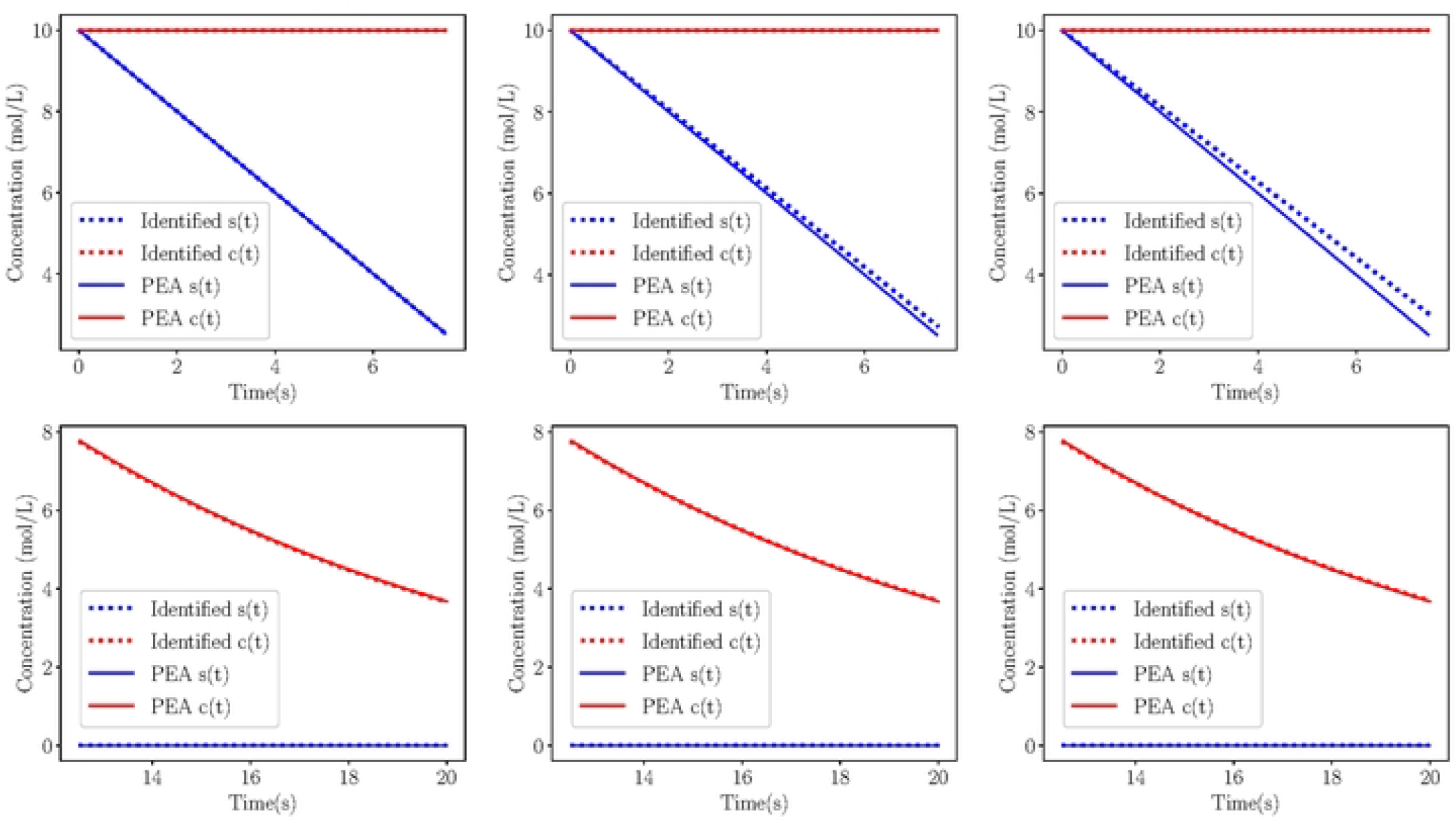
Comparisons. Solution comparison between the actual deterministic (solid) and identified from Weak SINDy (dotted) models of Michaelis-Menten simulations of Case 3 on the SIM, after spliting the data into regions where sQSSA (top) and rQSSA (bottom) models are valid. The data for the cases of no noise (left), additive (middle) and multiplicative noise (right) have been smoothened by the NODE. Parameters and initial conditions on Table 2.

A summary of the results from all stochastic simulations is displayed in Table 7, where the performance of Weak SINDy in identifying the full and reduced models is assessed and compared with that of the deterministic simulations. The deterministic models include results for both full and reduced models, while the stochastic models consider cases with additive and multiplicative noise at different noise levels (2% for additive and 1% for multiplicative). The results indicate that for the deterministic cases, Weak SINDy successfully identifies both the full and reduced models in Cases 1 and 2, whereas for Case 3, it fails to identify a reduced model when data from the reduced model is used. However, after partitioning the dataset into distinct regions (Case 3 Split 1 and Case 3 Split 2), corresponding to valid sQSSA and rQSSA models, Weak SINDy successfully identifies the correct reduced models in each sub-region.

**Table 7.**
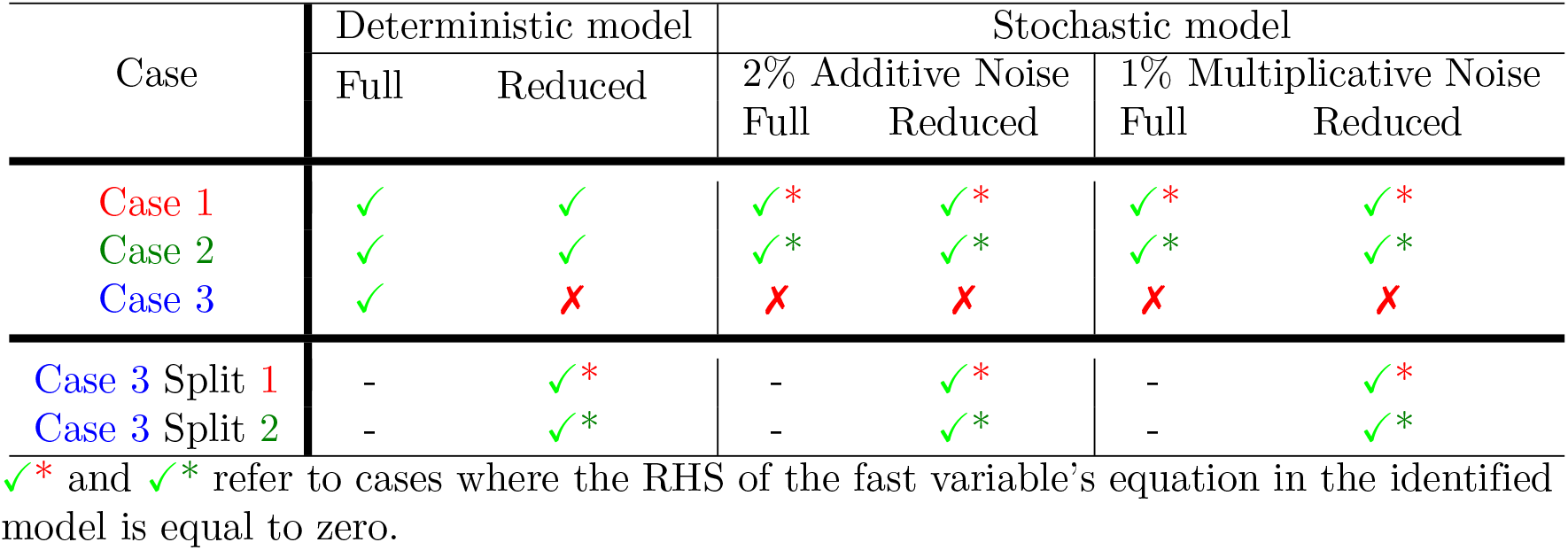
Performance of Weak SINDy on identifying the corresponding models of the data produced from the full and reduced deterministic and stochastic models.

## Discussion

We applied our proposed framework to the Michaelis-Menten system, a canonical model that captures the nonlinear and multi-scale nature of biological dynamics. Despite its simplicity, the model offers three analytically tractable time-scale regimes, namely sQSSA, rQSSA and PEA, serving as ideal test cases for validating data-driven methods. Through deterministic and stochastic simulations, we demonstrated that Weak SINDy can identify valid reduced models under appropriate conditions. However, it fails when applied to full datasets exhibiting transitions between regimes or containing non-standard nonlinearities. By integrating NODE-based vector field reconstruction, NN-based Jacobian estimation and CSP analysis, the framework successfully partitioned the data into dynamically consistent subregions, enabling accurate model identification across all cases, including noisy ones.

The summary of all simulations, shown in Table 7, confirms the strengths and limitations of Weak SINDy under different conditions. For deterministic data, the method recovered both full and reduced dynamics in Cases 1 and 2 but failed to reconstruct the PEA reduced model in Case 3. Upon partitioning the Case 3 dataset into separate sQSSA and rQSSA regimes, Weak SINDy correctly identified the reduced dynamics. This success persisted under noisy conditions, provided that the data were preprocessed with NODE. In cases where the fast variable’s derivative was near-zero, negative *R*^2^ values appeared. As previously discussed, such values should not be interpreted as failures but rather as artifacts of the QSSA structure and the limitations of variance-based metrics.

One limitation of the proposed method is its sensitivity to data quality. High noise levels or poor signal-to-noise ratios can obscure the underlying dynamics, particularly in systems governed by multiple time scales. Multiplicative noise, in particular, leads to significant distortion at high variable magnitudes, hampering the accuracy of the vector field and Jacobian estimation. While NODE can smooth noisy data to an extent, its effectiveness depends on the quality and density of the input data.

The performance of the NN components, including NODE and the Jacobian estimator, also hinges on careful tuning of hyperparameters. The number of nearest neighbors, architecture, learning rates, and training epochs all influence the accuracy of the estimated Jacobian. Incorrect configurations may lead to poor identification of fast and slow directions, undermining the CSP analysis. While our framework performed robustly under controlled conditions, real-world applications will likely require additional strategies for hyperparameter optimization and model validation.

Data sparsity presents another significant challenge. Sparse or short trajectories may not provide sufficient information for reliable vector field reconstruction, even with NODE. In such cases, preprocessing steps such as interpolation may be necessary to increase the sampling density. However, over-interpolation may introduce artifacts, necessitating a balance between data augmentation and preservation of the original dynamics.

To overcome sparsity and enhance model identification in limited-data settings, we propose incorporating Dynamic Mode Decomposition (DMD). This method can generate synthetic data from the latent space, enriching the available trajectory without requiring additional measurements. By expanding the data coverage, DMD supports more accurate vector field reconstruction and Jacobian estimation, thereby improving the reliability of the subsequent CSP and SINDy analyses. Also the DMD can serve as a comparison for the identified model predictions, when the ground truth is unkwown.

Extending the proposed framework to high-dimensional systems represents a critical direction for future research. Although CSP is inherently dimension-agnostic and theoretically remains applicable, the scalability of neural estimators poses practical challenges, as both computational cost and data requirements increase with dimensionality. Adapting the framework to real-world, high-dimensional datasets—such as those arising in systems biology, neuroscience, or clinical monitoring—will demand advances in both computational efficiency and data utilization. Given the complexity of biomedical systems, it is essential to develop integrative approaches that unify machine learning with mechanistic modeling [57–61]. Our method contributes to this goal, offering a foundation for scalable, data-driven multiscale analysis. Incorporating physics-informed constraints or domain-specific priors into the neural architecture offers a promising strategy to enhance generalization, reduce data demands, and broaden applicability across complex biological domains.

## Supporting information

### S1 Appendix

#### CSP Methodology

Assume that we have an *N*-dim. biological system, the evolution of which is described by the system of ODEs

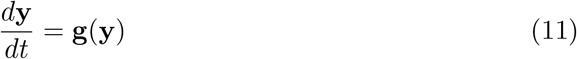

where **y** = [*y, y*, …, *y*]^*⊤*^ and 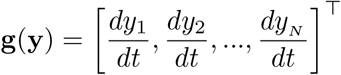 are the *N*-dim. Column state vector and vector field. According to CSP, any multi-scale system in the form of Eq. (11) can be cast into a form where the vector field **g**(**y**) is decomposed into *N* different terms that are defined by the CSP basis vectors **a**_*n*_ that span the fast and slow subspaces of the tangent space [35, 62]:

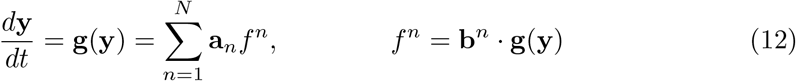

where each **a**_*n*_*f* ^*n*^ term is called a CSP mode and is a linear combination of all processes and variables of the system. **a**_*n*_ = **a**_*n*_(**y**) and **b**^*n*^ = **b**^*n*^(**y**) are the *N*-dim. CSP column and row, respectively, basis vectors of the *n*-th mode, and *f* ^*n*^ = *f* ^*n*^(**y**) is the amplitude of the *n*-th mode. The CSP basis vectors satisfy the orthogonality conditions 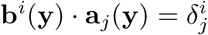[35, 63], *and proper adjustment of their signs sets f* ^*n*^ always positive. Each CSP mode describes subprocesses that act in different time scales and have different impact in the vector field. In particular each CSP basis vector **a**_*n*_ provides the direction along which a subprocess is evolving according to the *n*-th fastest time scale, and the amplitude *f* ^*n*^ measures how much the vector field is projected along the **a**_*n*_ direction; i.e. it measures the impact of the *n*-th CSP mode on **g**(**y**). The basis vectors that construct order by order the expressions in Eq. (12) are provided by CSP in an algorithmic manner [53, 54].

In the absence of the expolicit expression of the equations of the system in (11), the calculation of the CSP basis vectors is not always possible. Instead, approximations of the CSP basis vectors can be used to identify the fast/slow directions. Assuming that we have a dataset of *N* variables *y*_*i*_ (*i* = 1, …, *N*), the derivatives **g**(**y**) of which are calculated using finite differences, the Jacobian Matrix **J** of the system can be then estimated using the combination of NODE [33] and the NN of [34] described previously. Then, we can use the right, ***α***_***n***_, and left, ***β***^***n***^, eigenvectors of the calculated Jacobian matrix as a leading order approximation of the CSP basis vectors to span the fast and slow subspace of the vector field [35, 62]; **a**_*n*_ ≈ ***α***_*n*_ and **b**^*n*^ ≈ ***β***^*n*^. The time scales of the system in Eq. (11) can be approximated by the inverse modulo of the eigenvalues of **J**:

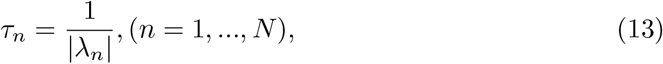

where *λ*_*n*_ are the non-zero eigenvalues of **J**. A zero eigenvalue refers to processes of infinite time scale. A negative (possitive) real part of the eigenvalue indicates a time scale of dissipative (explosive) nature.

Due to its nature, the system is expected to be evolving in different time scales, and it is assumed that the fastest of them, say *M*, are [i] of dissipative nature (i.e., the components of the system that generate them tend to drive the system towards equilibrium) and [ii] much faster than the rest, then the vector field **g**(**y**) can be decomposed into its fast and slow component:

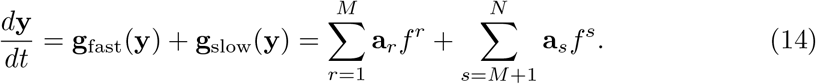

The fast **g**_fast_ (**y**) and slow **g**_slow_(**y**) components relate to the *M* fast and *N* −*M* slow, respectively, time scales. After the fast dissipative time scales act, they become exhausted and their related modes have no impact on the evolution of the system; i,e. the corresponding amplitudes become negligibly small. Then, a reduced model can be constructed as:

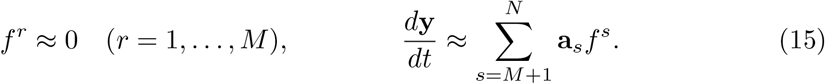

The left relation describes an *M*-dim. system of algebraic equations that define a low dimensional surface in the phase-space, called the *Slow Invariant Manifold (SIM)*, where the system is confined to evolve under the influence of the *M* fast exhausted time scales. These constraints refer to forming equilibria among fast reactions of the fast variables, and, thus, are characterized by the *M* fastest timescales, which are inherently dissipative in nature. The right relation is an *N*-dim. system of ODEs that governs the slow evolution of the system on the *SIM*, the dynamics of which are determined either by the fastest among the *N*− *M* slow, dissipative timescales or by explosive timescales, when the latter are present. The explosive timescales, in particular, are the time scales that tend to drive the system away from equilibrium. Such cases arise in all combustion systems [46, 64–66], but also in oscilating biological systems [44], population dynamics [67] and in systems describing cancer evolution [68].

#### CSP Pointer

Among the several developed algorithmic tools in the context of CSP, the CSP Pointer (Po) is an index that identifies the variables related the most to the a CSP mode. Through iterative analysis, Po assigns a numerical value to each variable and the CSP modes, reflecting its contribution to the fast or slow dynamics of the system [47]. The relation of the the *i*-th variable to the *m*-th CSP mode (*m* = 1, …, *M*) is calculated as:

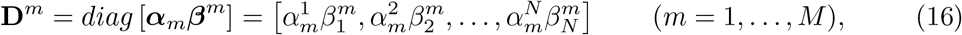

where, due to the orthogonality condition 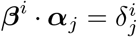, the sum of all *N* elements of **D**^*m*^ equals unity, i.e.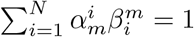[63, 69, 70]. The geometrical interpretation of the Po is presented in [49].

Each variable is associated differently to each CSP mode; A relatively large value of 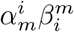 indicates that the *i*-th variable is strongly associated to the *m*-th CSP mode and the *m*-th timescale. For fast exhausted modes, the CSP Pointer identifies the variables related to fast dynamics; i.e. fast variables. In addition, a value of 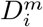 close to unity suggests that the *i*-th variable is in a quasi-steady-state [70].

The CSP Pointer tool is a critical extension of the CSP method, designed to systematically identify fast variables and processes within stiff systems. This allows us to quantitatively identify the dominant fast variables, facilitating the subsequent application of model reduction techniques such as Quasi-Steady State Approximation (QSSA) and Partial Equilibrium Approximation (PEA) [31, 70]. By using the CSP Pointer tool, it is possible to localize the regions in the system where model reduction can be applied most effectively, providing a powerful means of simplifying complex biological or chemical systems while retaining the essential dynamical features.

### S2 Appendix

#### CSP diagnostics of the deterministic models

Figure 9 (left) presents the developing time scales of the stochastic models of Cases 1 (top), 2 (middle) and 3 (bottom). A time scale gap is evident in all cases, indicative of the multi-scale character of the model. Similarly, the CSP amplitudes of the three cases are presented in the center column of the figure. It is shown that in Cases 1 and 2 the amplitude of the fastest dissipative mode becomes negligible, indicating the fast mode exhaustion and the forming equilibria of Eq. (15), based on which the reduced model can be constructed. Note that this does not hold in Case 3, where the initially vanishing amplitude is reactivated around *t* = 10*s*, before it becomes negligible again.

**Fig 9.**
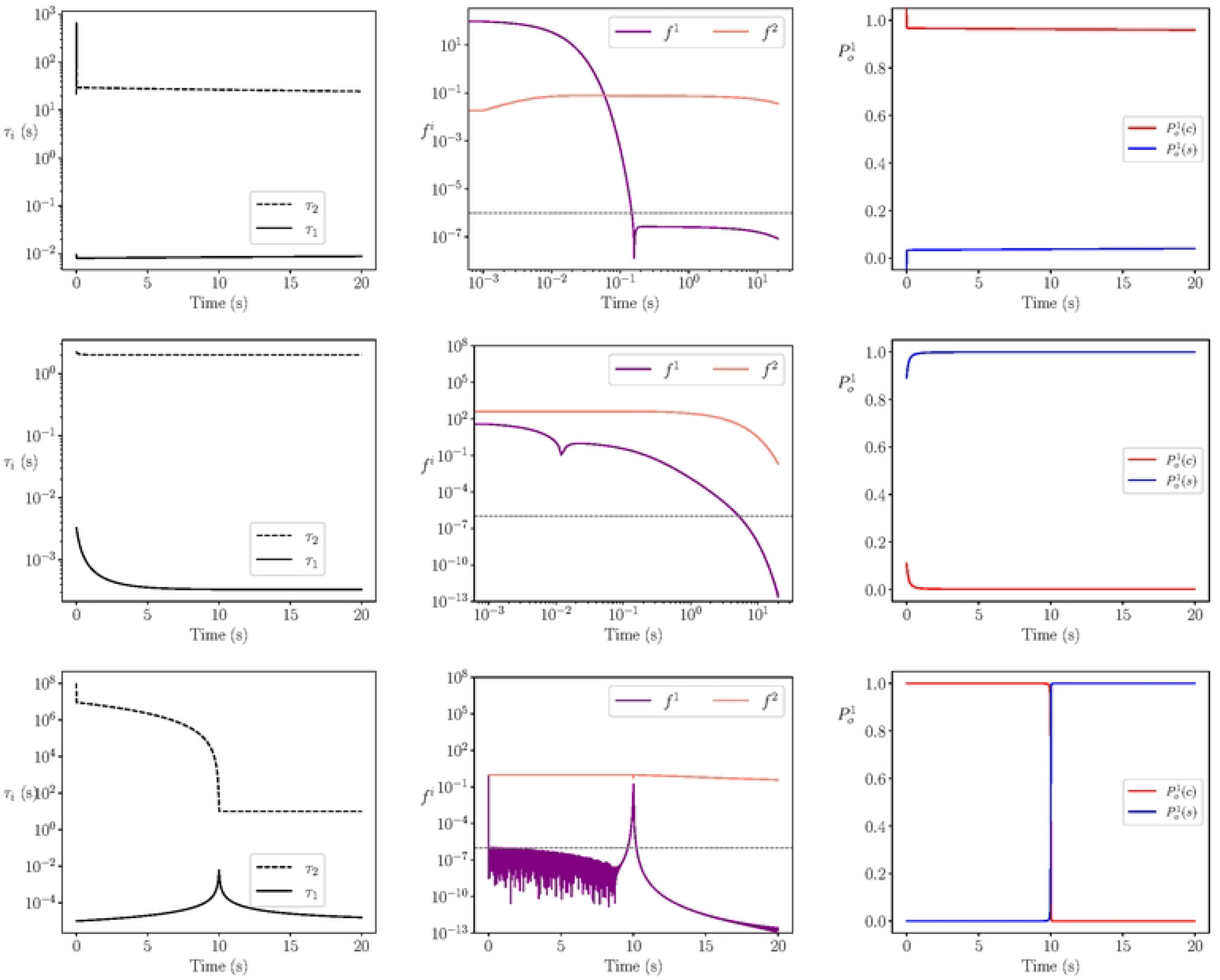
CSP Analysis. The evolution in time of the developing time scales (left), the amplitudes of the related CSP modes (center) and the CSP Pointer (right) for Case 1 (top), Case 2 (middle) and Case 3 (bottom).

The CSP Pointer (right) was used to identify the variables related the most to the fast dynamics througout the simulations. In Cases 1 and 2, the complex and substrate are identified to be the fast variables, respectively, indicating that valid sQSSA and rQSSA can be applied in each case. In Case 3, a shift in the dynamics is identified during the reactivation of the fast mode; the complex becomes the fast variable in place of the substrate. This shift corresponds to the transition between the regions of valid sQSSA and rQSSA (Figs 3 and 5).

### S3 Appendix

#### CSP diagnostics of the stochastic model of Case 3

Figure 10 (left) presents the developing time scales of the stochastic models (middle and bottom) in comparison to the reference case of no noise (top), using a reconstructed and smothened vector field from NODE. In all cases, a time scale gap is evident before and after the transition, with the fastest and driving time scale to be of the same order. The reactivation of the related to the fast mode amplitude (center) is captures in all cases on the transition, around *t* = 10*s*.

**Fig 10.**
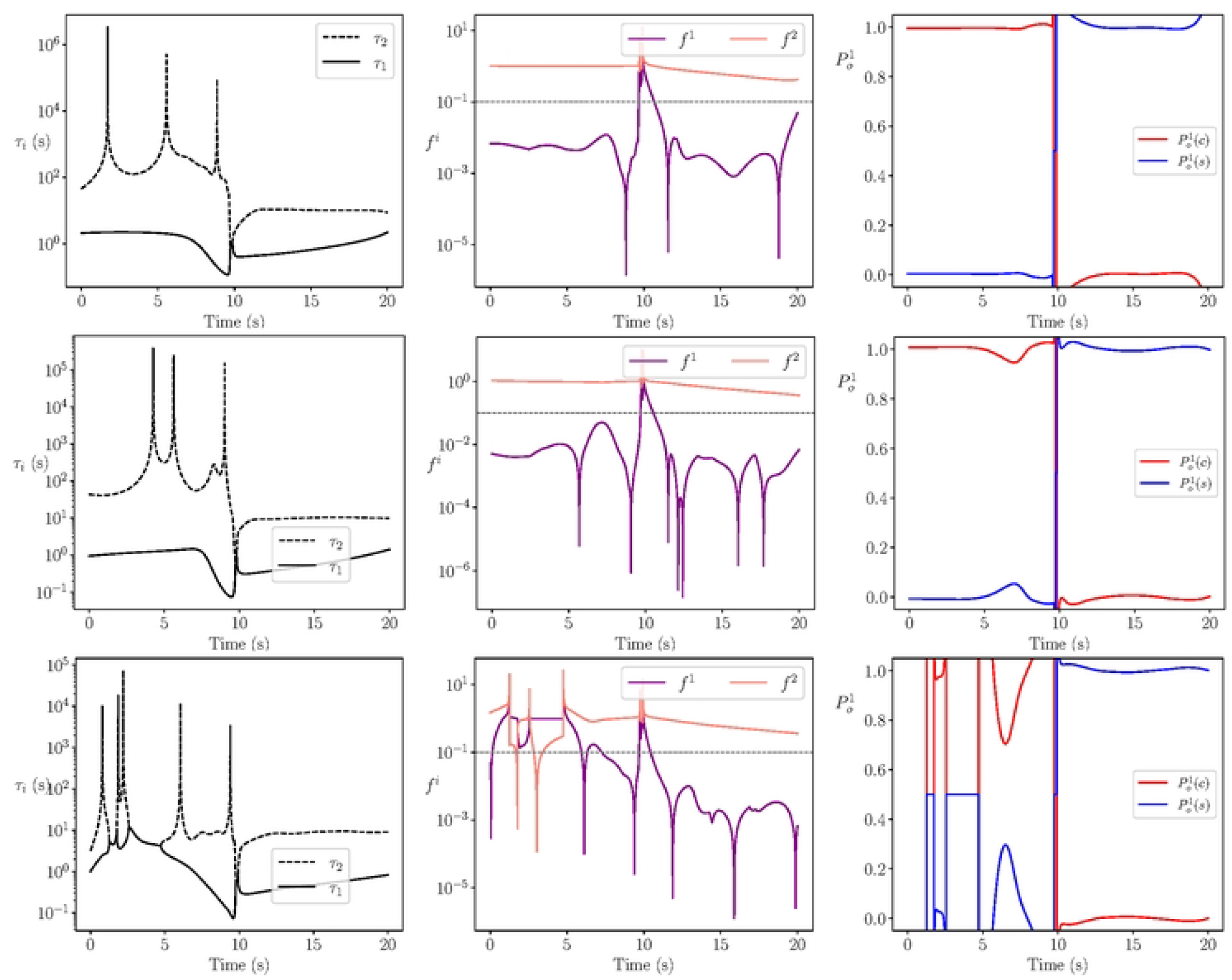
CSP Analysis. The evolution in time of the developing time scales (left), the amplitudes of the related CSP modes (center) and the CSP Pointer (right) as they were calculated from the reconstructed vector field from NODE for no noisy data (top), data with additive noise (middle) and data with multiplicative noise (bottom) of Case 3.

The CSP Pointer (right) was able to capture the shift in the dynamics during the reactivation of the fast mode, by identifying the fast variable of each region, indicative of valid sQSSA and rQSSA (Figs 3 and 5). These results demonstrate the robustness of our framework in handling noisy data, as the key outcomes remain consistent across different noise levels, enabling a reliable and unified analysis regardless of noise perturbations.

## Acknowledgments

Haralampos Hatzikirou would like to acknowledge the support of the Volkswagen Stiftung for the “Life?” initiative (grant number 96732). Haralampos Hatzikirou acknowledges the support of the RIG-2023-051 grant from Khalifa University. Haralampos Hatzikirou and Dimitris M. Manias would like to thank the UAE-NIH Collaborative Research grant AJF-NIH-25-KU.

## Author contributions

**Conceptualization:** Haralampos Hatzikirou

**Data curation:** Dimitris M. Manias, Ismaila Muhammed

**Formal analysis:** Ismaila Muhammed, Dimitris M. Manias, Haralampos Hatzikirou, Dimitris A. Goussis

**Funding acquisition:** Haralampos Hatzikirou

**Project administration:** Haralampos Hatzikirou

**Investigation:** Ismaila Muhammed, Dimitris M. Manias, Haralampos Hatzikirou, Dimitris A. Goussis

**Methodology:** Haralampos Hatzikirou, Dimitris A. Goussis

**Software:** Dimitris M. Manias, Ismaila Muhammed

**Supervision:** Haralampos Hatzikirou

**Validation:** Dimitris A. Goussis

**Visualization:** Dimitris M. Manias, Ismaila Muhammed

**Writing - original draft:** Ismaila Muhammed, Dimitris M. Manias

**Writing - review & editing:** Haralampos Hatzikirou, Dimitris A. Goussis

## Footnotes

1 Regarding the transition period (white region in Figs. 3 and 5), where no valid QSSA model can be constructed but only a PEA model, Weak SINDy identified the PEA model only when the nonlinear terms in the model were added in the library.

